# Extracellular lipids of *Camelina sativa*: Characterization of cutin and suberin reveals typical polyester monomers and novel functionalized dicarboxylic fatty acids

**DOI:** 10.1101/2020.06.21.163436

**Authors:** Fakhria M. Razeq, Dylan K. Kosma, Débora França, Owen Rowland, Isabel Molina

**Affiliations:** Department of Biology, Algoma University, Sault Ste. Marie, Ontario, Canada; Department of Biology and Institute of Biochemistry, Carleton University, Ottawa, Ontario, Canada; Department of Biochemistry and Molecular Biology, University of Nevada, Reno, Nevada 89557, USA

**Keywords:** *Camelina sativa*, lipid polyesters, suberin, cutin, aliphatic and aromatic monomers, GC-MS, GC-FID, TEM

## Abstract

*Camelina sativa* is relatively drought tolerant and requires less fertilizer than other oilseed crops. Various lipid- and phenolic-based extracellular barriers of plants help to protect them against biotic and abiotic stresses. These barriers, which consist of solvent-insoluble polymeric frameworks and solvent-extractable waxes, include the cuticle of aerial plant surfaces and suberized cell walls found, for example, in periderms and seed coat. Cutin, the polymeric matrix of the cuticle, and the aliphatic domain of suberin are fatty acid- and glycerol-based polyesters. These polyesters were investigated by base-catalyzed transesterification of *C. sativa* aerial and underground delipidated tissues followed by gas chromatographic analysis of the released monomer mixtures. Seed coat and root suberin had similar compositions, with 18-hydroxyoctadecenoic and 1,18-octadecenedioic fatty acids being the dominant species. Root suberin presented a typical lamellar ultrastructure, but seed coats showed almost imperceptible, faint dark bands. Leaf and stem lipid polyesters were composed of fatty acids (FA), dicarboxylic acids (DCA), ω-hydroxy fatty acids (OHFA) and hydroxycinnamic acid derivatives (HCA). Dihydroxypalmitate (DHP) and caffeic acid were the major constituents of leaf cutin, whereas stem cutin presented similar molar proportions in several monomers across the four classes. Unlike the leaf cuticle, the *C. sativa* stem cuticle presented lamellar structure by transmission electron microscopy. Flower cutin was dominated by DHP and did not contain aromatics. We found striking differences between the lipid polyester monomer compositions of aerial tissues of *C. sativa* and that of its close relatives *Arabidopsis thaliana* and *Brassica napus*.

**Graphical Abstract:** 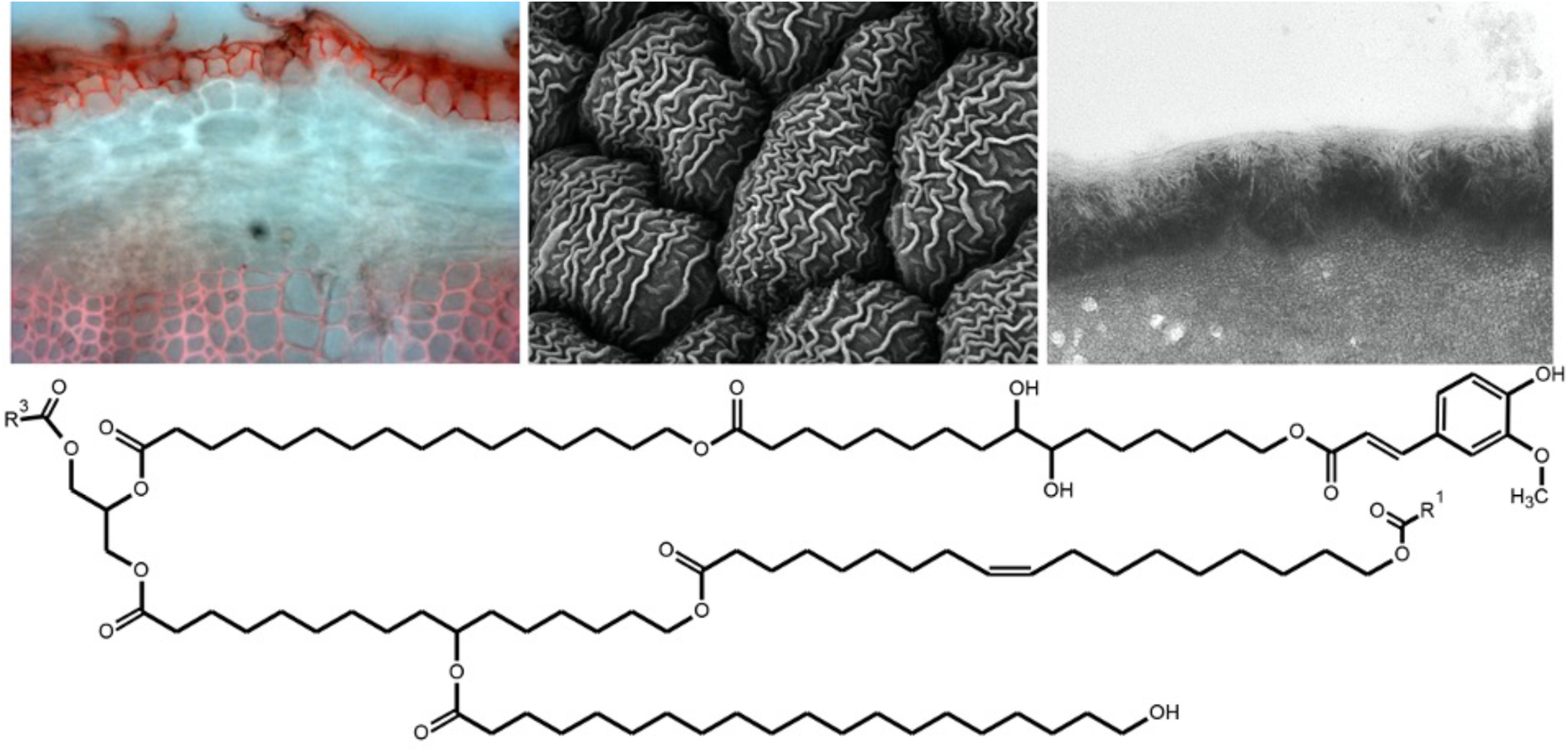

*Camelina sativa* leaf and stem cutin is dominated by 16:0 dihydroxy fatty acid and aromatics, with dicarboxylic fatty acids representing 20-30 % of the monomers. Suberin of root and seed coat is largely composed of 18:1 dicarboxylic and ω-hydroxy fatty acids.

**Highlight bullet points:**

- *Camelina sativa* is an oil crop tolerant to biotic and abiotic stresses
- Extracellular lipid polyesters may in part confer these attributes
- Dihydroxypalmitate and caffeic acid were major components of *C. sativa* leaf cutin
- Flower cutin lacked aromatics and contained monomers not previously reported
- Root and seed coat suberin was dominated by 18:1 ω-hydroxy and dicarboxylic fatty acids
- C18 monounsaturated photo-oxidation products were found in leaf cutin and suberin

## 1. Introduction

Plant extracellular lipids form physical barriers that control water and solute movement as well as gas exchange. Because of their barrier properties, extracellular lipids are critical to protect plants against various biotic and abiotic stresses, such as drought, salinity and attack by pathogenic microorganisms. These surface lipids are organized into complex hydrophobic barriers of polymerized polyesters and associated waxes, and include: cutin found in the cell walls of epidermis of all aerial tissues, and suberin found in the cell walls of various internal and external tissue layers, such as root endodermis and periderm, and seed coat (Kolattukudy, 1981). The lipid composition of both cuticle and suberin varies considerably between plants and can be altered by environmental factors (Franke et al., 2009; Jetter et al., 2008; Kosma et al., 2009; Kosma and Jenks, 2007; Pollard et al., 2008).

Cutin is a polymer composed of aliphatics (typically C16 and C18 ω-hydroxy, polyhydroxy, epoxy and α,ω-dicarboxylic fatty acids), glycerol and usually small amounts of phenolic compounds (Beisson et al., 2012; Nawrath, 2006). The chemical composition, structure and thickness of cutin are highly variable depending on the plant species and the tissue types. For example, leaf and stem cutin of *Arabidopsis thaliana* contain unusually high amounts of C16 and C18 α,ω-dicarboxylic fatty acids, whereas Arabidopsis flower cutin mainly contains poly-hydroxy fatty acids (Beisson et al., 2012).

Suberin is chemically similar to cutin, but with higher phenolic and C20-C24 aliphatic content (Vishwanath et al., 2015). Suberin consists of polyphenolic and polyaliphatic domains linked together in a three dimensional polyester network, which often has the appearance of alternating light and dark bands when viewed by transmission electron microscopy, collectively referred to as suberin lamellae (Pollard et al., 2008). The polyphenolic part consists largely of hydroxycinnamic acids (mostly ferulic acid, but also *p*-coumaric and caffeic acids), and small amounts of monolignols (*p*-coumaryl, coniferyl and sinapyl alcohols). The polyaliphatic part of suberin mainly consists of glycerol, long-chain (C16 and C18) and very-long-chain (>C18) α,ω-dicarboxylic fatty acids, ω-hydroxy fatty acids, mid-chain-oxidized fatty acids, primary fatty alcohols (usually 18:0-22:0), and very-long-chain fatty acids (typically 22:0 and 24:0) (Beisson et al., 2012; Graça and Santos, 2007; Ranathunge et al., 2011). As with cutin, suberin composition varies within and between plant species, especially in the relative proportions of the constituent monomers (Holloway, 1983).

Cutin is deposited on the outermost surfaces of epidermal cell walls of all aerial organs of the plant (Beisson et al., 2012). Suberin is deposited on the inner face of the primary cell wall of various internal and external cellular tissue layers, such as endodermis of young roots, periderms of mature roots and tubers, seed coat integuments, bark tissue, abscission zones, and cotton fibers (Bernards, 2002; Ranathunge et al., 2011). Suberin is also deposited in response to wounding and pathogenic attack (Lulai and Corsini, 1998). Both cutin and suberin, as well as their associated waxes, are involved in controlling solute and water transport across cell walls and provide a barrier against pathogens (Pollard et al., 2008). In many plant species, a non-depolymerizable fraction known as cutan remains after ester-bound monomers are extracted. This polymer may be composed of ether-bond unsaturated aliphatics, aromatics and polysaccharides (Leide et al., 2020; Nip et al., 1986; Villena et al., 1999). A similar polymer, termed suberan, has been found associated with suberized tissues (Tegelaar et al., 1995; Turner et al., 2013).

Recent genetic and biochemical studies in model plants, such as *Arabidopsis thaliana*, have improved our understanding of the biosynthesis, regulation, transport and deposition of cutin and suberin (reviewed in Cohen et al., 2017; Fich et al., 2016; Li-Beisson et al., 2016; Schreiber, 2010; Vishwanath et al., 2015). Studies in other model plants, such as *Solanum lycopersicum* (tomato) and *Solanum tuberosum* (potato), have provided detailed knowledge about polyester structure and biosynthesis for cutin and suberin, respectively (Graça and Pereira, 2000; Isaacson et al., 2009; Matas et al., 2011; Schreiber et al., 2005).

Detailed lipid polyester profiles have been described for Arabidopsis flowers (Beisson et al., 2007; Li-Beisson et al., 2009), leaves (Bonaventure et al., 2004; Franke et al., 2005), stems (Bonaventure et al., 2004; Suh et al., 2005), seeds (Beisson et al., 2007; Molina et al., 2008, 2006), and roots (Beisson et al., 2007; Franke et al., 2005). A thorough characterization of the protective surface lipids of wild-type plant species provides a valuable reference to which polyester compositions of mutants can be compared to. This approach may help to unravel the complexity of cutin and suberin chemical composition, and to understand the functions of associated genes by reverse genetics.

*Camelina sativa* (L.) Crantz is an emerging oilseed crop with oil content rich in omega-3 fatty acids and with composition suitable for biofuel or industrial oil production, while its seed cake provides high nutritional value for animal feed and could be used for production of various biodegradable materials (Faure and Tepfer, 2016; Obour K, 2015). *C. sativa* is relatively drought and frost tolerant and requires less fertilizer than other oilseed crops, such as canola and soybean. In addition, *C. sativa* appears to be resistant to many pests and pathogens that affect other oilseed crops, such as blackleg disease and striped flea beetle (Soroka et al., 2015; Vollmann and Eynck, 2015). High natural genetic diversity and its short growth cycle provides the opportunity to create new plant varieties through conventional plant breeding or genetically engineered varieties (Lu and Kang, 2008), making *C. sativa* ideal for both research and field applications (Bansal and Durrett, 2016).

A detailed description of the protective surface lipids of *C. sativa* may provide insights into its drought-tolerant and pathogen-resistant properties, and also provide an additional source of high-value lipid components that can be extracted from the plant. In this study, the chemical compositions and ultrastructural features of extracellular lipids extracted from aerial and subterranean tissues of *C. sativa* were analyzed using gas chromatography and microscopy. This report, together with our previous study detailing the chemical composition of *C. sativa*’s extracellular waxes (Razeq et al., 2014), provide complete qualitative and quantitative information on the waxes and lipid polyesters extracted from *C. sativa* boundary tissues.

## 2. Results and Discussion

### 2.1. Amounts and ultrastructure of C. sativa extracellular lipid-based polymers

Cutin monomers were isolated after depolymerization by NaOMe-catalyzed methanolysis of solvent-extracted dry residues from whole leaf, stem and flower tissues (Jenkin and Molina, 2015; Molina et al., 2006). Quantitative analyses by GC-FID, using internal standards, showed concentrations of ca. 2.2, 0.8 and 1.8 mg·g^-1^ DW of total identified monomers in leaf, stem and flower, respectively (**Table 1**). There is an apparent discrepancy with one published report on *C. sativa* leaf cutin, where the cutin concentration per leaf area unit was four times larger than the amount reported here (Tomasi et al., 2017). However, that difference is mostly because of the large proportions of fatty acids derived from membranes, namely unsaturated fatty acids and 2-hydroxy fatty acids, which were thoroughly removed in our preparations. By comparison, the closely related species *Arabidopsis thaliana* had cutin monomer loads that were half and two thirds of the amounts found in *C. sativa* stem and flower depolymerizates, respectively (Li-Beisson et al., 2013), whereas the load of leaf cutin in Arabidopsis was comparable to that of *C. sativa* (Franke et al., 2005). However, *C. sativa* leaf adaxial cuticles determined by transmission electron microscopy (TEM) were about 70 nm-thick (**Figure 1A**), which is more than two times thicker than *A. thaliana* cuticles (Franke et al., 2005). Whereas the cutin amounts in leaves of both species are similar, *Camelina sativa* leaf wax coverage is at least four times larger than that of *A. thaliana* leaves (Razeq et al, 2014); it is unclear whether differences in wax deposition between these species explains the differences observed in cuticle thickness. Abaxial *C. sativa* leaf cuticles were slightly thinner (50 nm) than adaxial cuticles and showed one or two electron-translucent lamellae (**Figure 1B**). The stem epidermis (**Figure 1C-D**) was covered with a much thicker cuticle (about 220 nm), correlating with a cutin amount per area unit that was more than 10 times higher than the cutin coverage in leaves (**Table 1**). Stem cuticles presented electron-translucent lamellae that were randomly oriented, reminiscent of those observed in cuticles of *Cuscuta gronovii* (Heide-Jørgensen, 1991; Jeffree, 1996). Differences in chemical composition between both cutin polymers could influence the ultrastructural arrangement (discussed below). It has been also suggested that the lamellar structure in some species can be attributed to alternating waxes and cutin, or to a combination of soluble and saponifiable lipids associated with cutan (reviewed by Jeffree, 1996). Flower parts were not analyzed by TEM, and petal cuticles observed by scanning transmission microscopy (SEM) presented characteristic nanoridge structures also observed in Arabidopsis petals (**Figure 1E-F**) (Li-Beisson et al., 2009).

**Table 1.** Total amounts of lipid polyester monomers isolated from solvent-extracted *Camelina sativa* stem, leaf, flower, seed and root residues by NaOMe-catalyzed transmethylation.

| Organ | mg g<sup>-1</sup> DW |  | µg dm<sup>-2</sup> |  |
| --- | --- | --- | --- | --- |
|  | Mean | SD | Mean | SD |
| Leaf<sup>1</sup> | 2.17 | 0.19 | 46.0 | 6.8 |
| Stem<sup>1</sup> | 0.77 | 0.19 | 526.0 | 128.0 |
| Flower<sup>2</sup> | 1.79 | 0.74 | n.d. |  |
| Root<sup>3</sup> | 9.10 | 1.67 | n.d. |  |
| Seed<sup>4</sup> | 2.73 | 0.34 | 1738.0 | 305.0 |

Data are mean with SD ( $n = 4$ ). DW = dry weight. n.d. = area not determined.
<sup>1</sup>For leaf and stem cutin analysis, samples were prepared from 8-week-old plants; leaves #14-17 and stem from the same area (~25 cm above the soil or 'middle stem' as defined in Razeq et al. 2014) were analyzed.
<sup>2</sup>Fully opened flowers were harvested for cutin determinations.
<sup>3</sup>Roots were harvested from 4-week-old plants grown on agar.
<sup>4</sup>Seeds were fully mature dry seeds.

**Figure 1.**
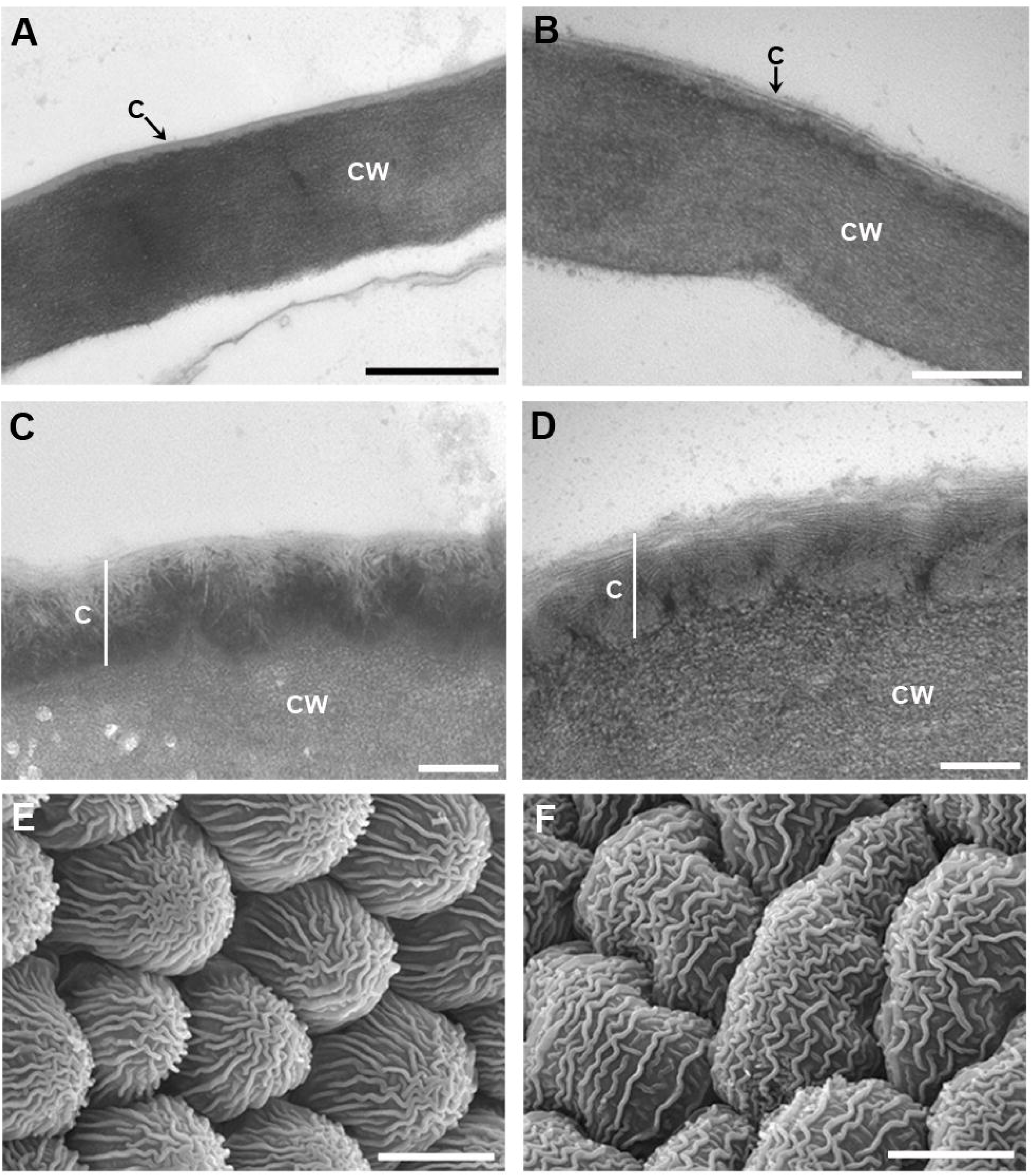
Ultrastructure of *Camelina sativa* cuticles. Transmission electron microscopy images of cross-sections of adaxial (A) and abaxial (B) leaves, and top (C) and bottom (D) stems. Scanning electron microscopy images of adaxial (E) and abaxial (F) petal surfaces. Scale bars: 500 nm (A), 200 nm (B, C, D), and 10 *µ*m (E, F). C, cuticle; CW, cell wall.

Suberized cell walls were identified in root periderm and seed coat sections (**Figure 2**). The periderm of 5-week-old roots had two layers of cells presenting characteristic red staining upon treatment with sudan red 7B (**Figure 2A**) as well as blue autofluorescence (**Figure 2B**). Observation by transmission electron microscopy revealed a lamellar ultrastructure in both the root endodermis (**Figure 2C**) and periderm (**Figure 2D**). Suberization was also evident on the palisade cell walls of seed coat sections (**Figure 2 E-F**), but in these tissues the darker bands of the lamella were almost imperceptible and reminiscent to the lamellae observed in the outer integument of Arabidopsis seed coats (Yadav et al., 2014). It is unknown whether differences in ultrastructure may result from different monomer chemistries or the arrangement of these units in the polymer. In particular, we did not find substantial differences between the chemical composition of the major *C. sativa* root and seed coat suberin ester-bound monomers (section 2.2). The total amounts of suberin monomers were 9.10 and 2.90 mg g^-1^ root and whole seed dry cell wall residues, respectively (**Table 1**). It should be noted that the seed suberin load includes a small contribution of cutin monomers from the embryo and internal seed coat cuticles (Molina et al., 2006). The root suberin monomer yield from *C. sativa* was comparable to the amount reported for Arabidopsis (7.2 mg g^-1^ DW; Li et al., 2007) whereas seeds contained about one third of the amount of lipid polyesters reported for Arabidopsis (8.6 mg g^-1^ seed residue; Molina et al., 2006). Differences in seed lipid polyesters that are normalized to total cell wall delipidated tissue may lead to inaccurately concluding that *C. sativa* seeds have lower suberin content than Arabidopsis. Given the smaller size of Arabidopsis seeds, it is expected that lipid polyesters, which are mostly localized to the seed coat, will be more concentrated in a preparation that contains more seeds -and thus higher seed surface area-in a given mass. In fact, when the polyester monomer amounts are normalized to surface area, *C. sativa* seeds contain ca. 33% higher loads than Arabidopsis (Table 1; Molina et al., 2006).

**Figure 2.**
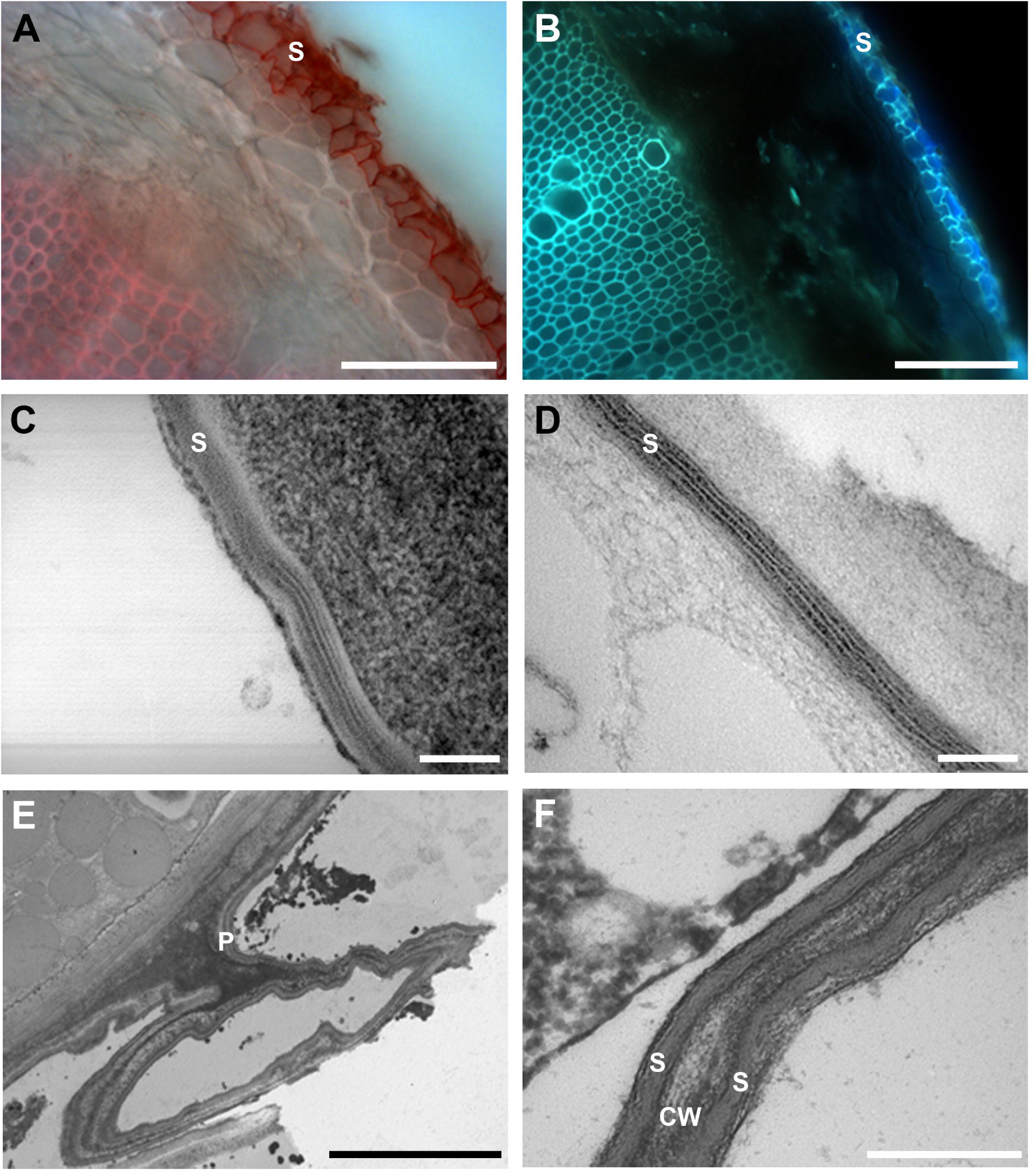
Suberin deposition in roots and seed coats of *Camelina sativa*. Root cross sections showing suberized root periderm stained with Sudan Red (A) or viewed via blue-yellow suberin autofluorescence (B). Transmission electron microscopy (TEM) image of root endodermis (C) and TEM image of root periderm (D). TEM image of seed coat showing suberized palisade cell walls (E, F). Scale bars: 100 *µ*m (A, B), 100 nm (C, D), 5 *µ*m (E), and 500 nm (F). CW, cell wall; P, palisade layer; S, suberin.

### 2.2. Characterization of C. sativa lipid polyesters

The sodium methoxide (NaOMe)-catalyzed methanolysis method employed in this study followed by solvent partition to recover monomers and their subsequent conversion to trimethylsilyl (TMSi) ether derivatives for GC-MS(EI) analysis, yields distinctive ions that allow for monomer identification. Upon transmethylation, ester-bound acids are converted to corresponding methyl esters and free hydroxyl acids. If epoxides are present in the polymer, the oxirane ring is opened during solvolysis in basic methanol giving a substitution product with methoxy and hydroxyl groups in adjacent carbons that is readily identifiable after derivatization (Holloway, 1974; Kolattukudy and Agrawal, 1974). Mid-chain substituted fatty acids are also commonly found in cutins. These monomers undergo α-cleavage on either side of the substituent (e.g. –CH[OTMSi]-) giving diagnostic mass spectra (Kolattukudy, 1984). *C. sativa* cutins depolymerized with this method yielded both typical, previously reported monomers, and monomers that were tentatively identified by their mass spectra (MS) and retention time and have not been described previously in the literature.

#### Leaf, stem and flower cutin

In *C. sativa* cutin from the three organs studied, 10,16-dihydroxy hexadecanotate methyl ester (or 10,16-dihydroxypalmitate; DHP) was either the predominant or a major component, representing 17, 10 and 47 % of the monomers from leaf, stem and flower tissues, respectively, with smaller proportions of the co-eluting 9-hydroxy positional isomer (**Table 2**). This monomer was identified by comparison to published mass spectra (Eglinton and Hunneman, 1968, Eglinton et al., 1968; Holloway, 1982; Holloway and Deas, 1971) and by its retention time on a nonpolar column (**Figure 3A**). In a given cutin, monomers are often classified according to the chain-length of the most predominant monomers belonging to the C_16_ or C_18_ families (Holloway, 1982). Despite the predominance of DHP, leaf or stem *C. sativa* cutin can be classified as mixed C_16_/C_18_ cutins, since the C_16_ functionalized monomers accounted for 28% and 17% of the leaf and stem monomers, respectively, whereas C_18_ and odd chain fatty acid derivatives altogether constituted 28% and 36 % of the released monomers from leaf and stem cutins, respectively (**Table 2**). Flowers, on the other hand, had a typical C_16_ cutin with 60% of the monomers corresponding to functionalized 16:0 fatty acids. The flower cutin monomer profile overlapped only partially with that of the leaf and stem cutin. For example, hydroxycinnamates and several minor in-chain substituted fatty acids were clearly absent from these tissues and some mid-chain hydroxylated fatty acid derivatives were exclusively found in flower cutin.

**Table 2.**
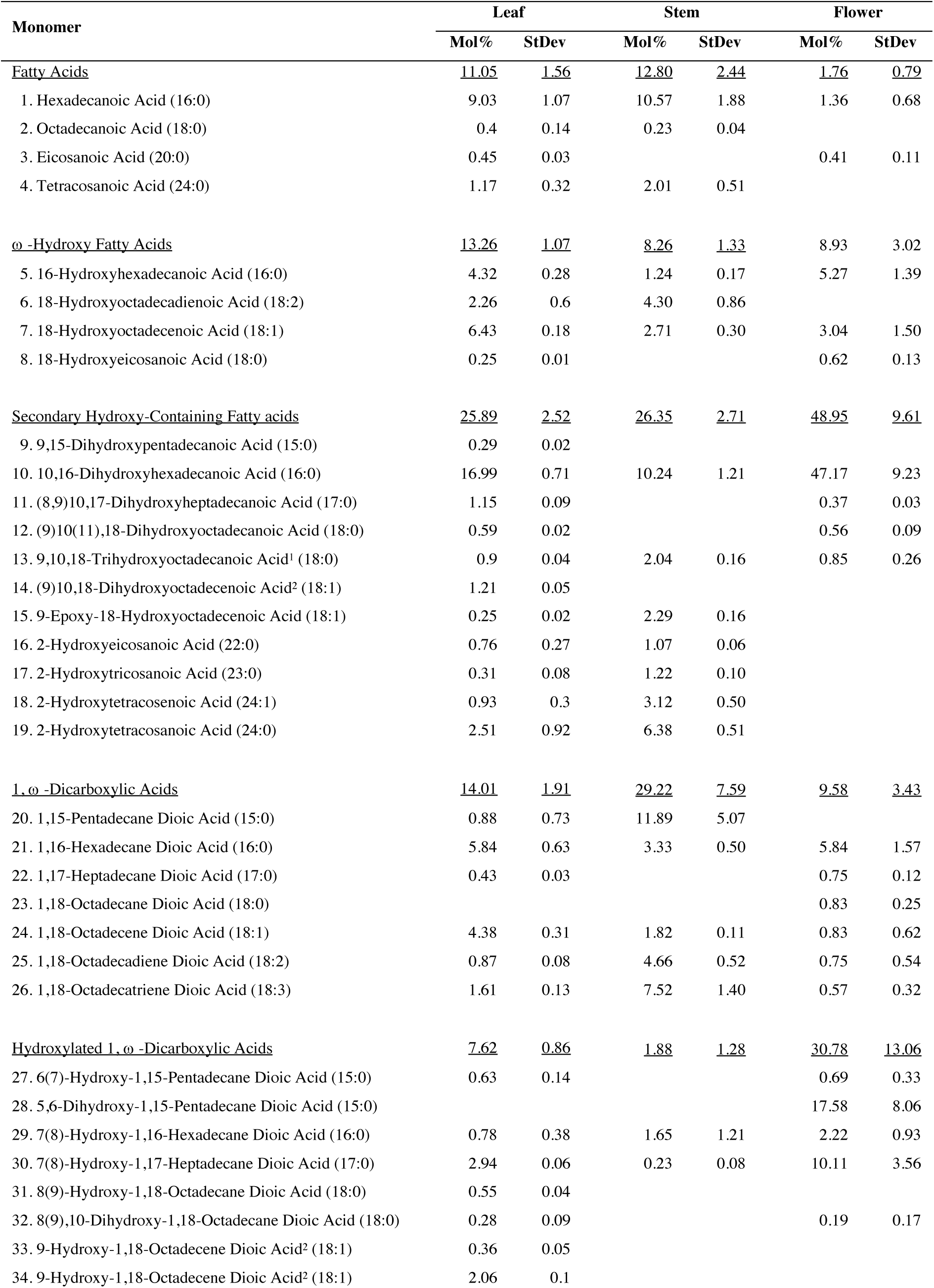
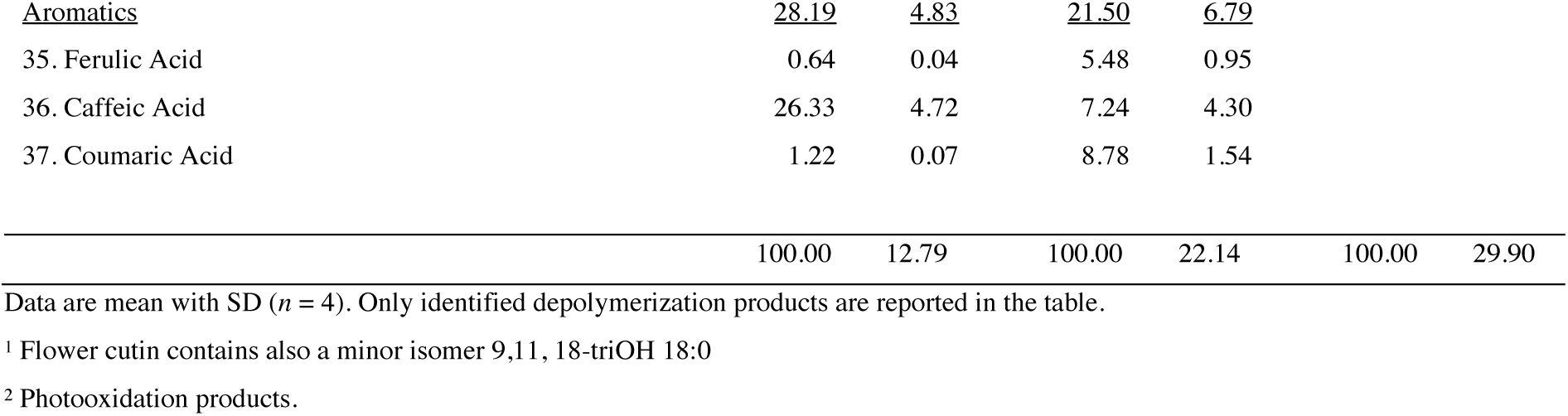
Detailed cutin monomer compositions of *C. sativa* leaf, stem and flower.

**Figure 3.**
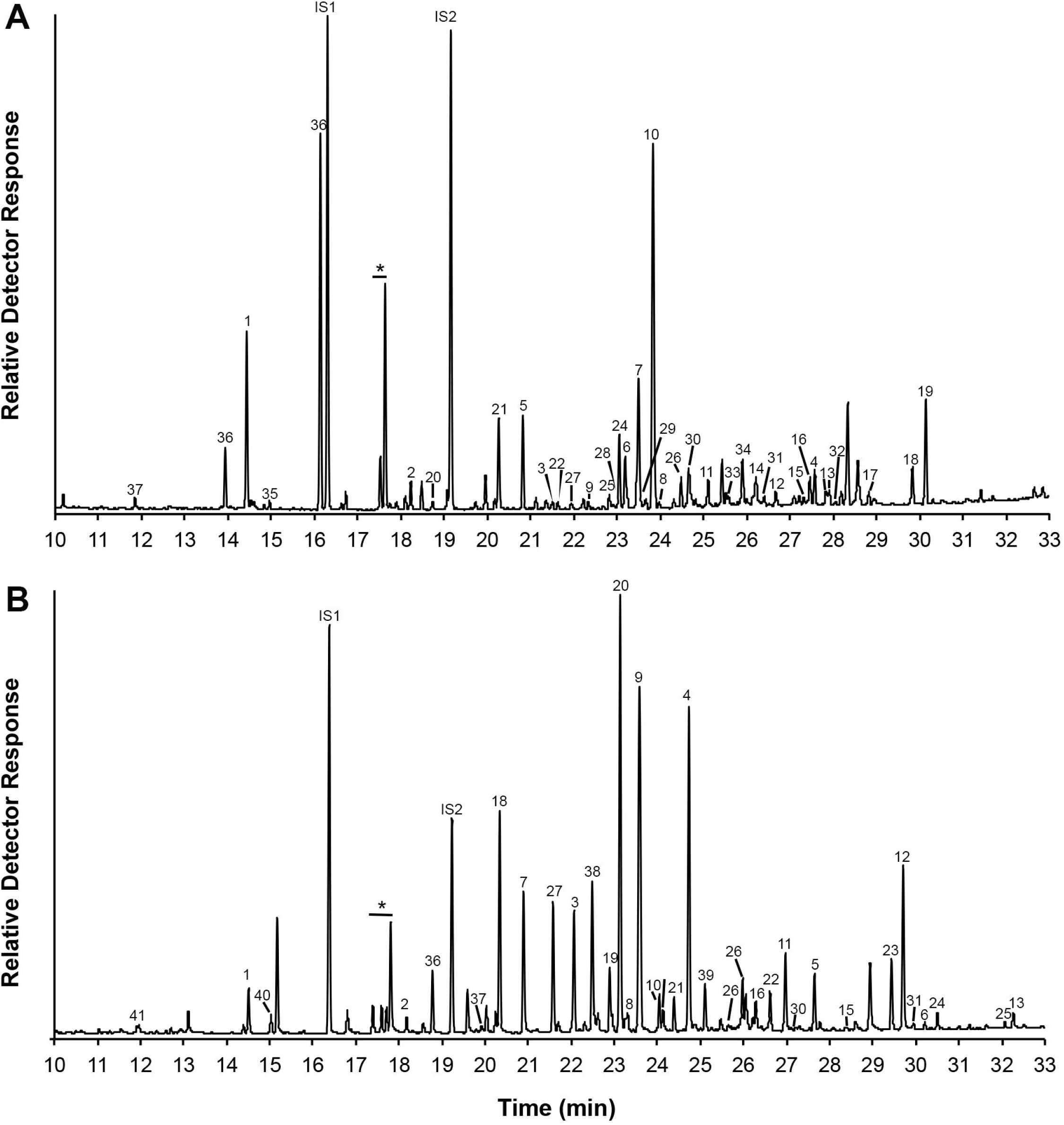
Annotated chromatograms of TMSi derivatives of *C. sativa* leaf cutin (A) and root suberin (B) monomers. Peak numbers correspond to monomers listed in Table 2 (cutin monomers) and Table 3 (suberin monomers). Internal standard (IS): 17:0 fatty acid methyl ester (IS1) and 15-hydroxy 15:0 fatty acid methyl ester (IS2). Asterisks indicate peaks of residual unsaturated fatty acids from membranes, not considered part of the polyester.

#### Root and seed coat suberin

Whole *C. sativa* mature seeds and 5-week-old roots were exhaustively delipidated and chemical depolymerization was carried out on cell wall-enriched dry residues (Molina et al., 2006). Seeds are complex organs with several tissues containing extracellular lipid polymers (Molina et al., 2008; De Giorgi et al., 2015), and the monomers identified and quantified on whole seeds may derive from one or more of such tissues. Most of the monomers released from *C. sativa* seeds can be classified as typical of suberin components, because 1) dimethyl octadecene-1,18-dioate and methyl 18-hydroxy-octadecenoate (both of which are low in cutins characterized in this study) were predominant in seed samples, and 2) the monomer profile was very similar to that of the root suberin samples (**Table 3**). Although dimethyl octadecene-1,18-dioate and methyl 18-hydroxy-octadecenoate are typical constituents of many suberins, including potato periderm (Kolattukudy and Dean, 1974), and root tissues of *Zea mays* and *Ricinus communis* (Schreiber et al., 2005), both monomers are also found in aerial cuticles of phylogenetically related species, namely Arabidopsis and *Brassica napus* (Bonaventure et al., 2004). Therefore, any classification as cutin or suberin solely based on chemical analyses should be taken cautiously. However, our microscopical analyses of root and seed sections confirmed the presence of suberin on cell walls of root periderm (**Figure 2 A,B,D**), root endodermis (**Figure 2C**), and seed coat palisade cells (**Figure 2 E-F**). As a result, both seed and root monomers are reported as suberin components in **Table1** and **Table 3**.

**Table 3.** Detailed suberin monomer compositions of *C. sativa* root periderm and seed coat.

| Monomer | Root |  | Seed |  |
| --- | --- | --- | --- | --- |
|  | Mol% | StDev | Mol% | StDev |
| <u>Fatty Acids</u> | <u>15.73</u> | <u>1.56</u> | <u>4.83</u> | <u>0.55</u> |
| 1. Hexadecanoic Acid (C16:0) | 1.08 | 0.15 | 2.69 | 0.32 |
| 2. Octadecanoic Acid (C18:0) | 0.48 | 0.30 | 1.31 | 0.14 |
| 3. Eicosanoic Acid (C20:0) | 2.34 | 0.14 | 0.17 | 0.04 |
| 4. Docosanoic Acid (C22:0) | 9.89 | 0.65 | 0.23 | 0.02 |
| 5. Tetracosanoic Acid (C24:0) | 1.69 | 0.16 | 0.44 | 0.04 |
| 6. Hexacosanoic Acid (C26:0) | 0.25 | 0.16 |  |  |
| <u>ω-Hydroxy Fatty Acids</u> | <u>39.04</u> | <u>3.74</u> | <u>28.763</u> | <u>2.62</u> |
| 7. 16-Hydroxyhexadecanoic Acid (C16:0) | 5.09 | 1.86 | 5.35 | 0.94 |
| 8. 18-Hydroxyoctadecadienoic Acid (C18:2) | 1.27 | 0.07 | 1.28 | 0.44 |
| 9. 18-Hydroxyoctadecenoic Acid (C18:1) | 20.02 | 0.98 | 18.18 | 0.95 |
| 10. 18-Hydroxyeicosanoic Acid (C18:0) | 1.63 | 0.16 |  |  |
| 11. 20-Hydroxyeicosanoic Acid (C20:0) | 3.66 | 0.22 | 0.86 | 0.21 |
| 12. 22-Hydroxydocosanoic Acid (C22:0) | 7.00 | 0.39 | 2.24 | 0.05 |
| 13. 24-Hydroxytetracosanoic Acid (C24:0) | 0.17 | 0.05 | 0.81 | 0.02 |
| <u>Secondary Hydroxy-Containing Species</u> | <u>0.39</u> | <u>0.16</u> | <u>3.79</u> | <u>0.51</u> |
| 14. 9,10,18-Trihydroxyoctadecenoic Acid (C18:1) |  |  | 1.82 | 0.17 |
| 15. 9,10,18-Trihydroxyoctadecanoic Acid (C18:0) | 0.04 | 0.05 | 0.65 | 0.06 |
| 16. (9)10,18-Dihydroxyoctadecenoic Acid<sup>1</sup> (C18:1) | 0.14 | 0.03 | 0.80 | 0.14 |
| 17. 21-Hydroxydocosanoic Acid (C22:0) |  |  | 0.51 | 0.13 |
| 18. 2-Hydroxytetracosanoic Acid (24:0) | 0.21 | 0.08 |  |  |
| <u>1,ω-Dicarboxylic Acids</u> | <u>30.86</u> | <u>1.97</u> | <u>34.65</u> | <u>1.64</u> |
| 19. 1,16-Hexadecane Dioic Acid (C16:0) | 8.71 | 0.42 | 8.56 | 0.27 |
| 20. 1,18-Octadecadiene Dioic Acid (C18:2) | 2.44 | 0.12 | 1.26 | 0.08 |
| 21. 1,18-Octadecene Dioic Acid (C18:1) | 15.92 | 1.13 | 19.79 | 1.06 |
| 22. 1,18-Octadecatriene Dioic Acid (C18:3) | 0.14 | 0.01 |  |  |
| 23. 1,20-Eicosane Dioic Acid (C20:0) | 1.31 | 0.09 | 0.16 | 0.01 |
| 24. 1,22-Docosane Dioic Acid (C22:0) | 2.28 | 0.14 | 2.87 | 0.09 |
| 25. 1,23-Tricosane Dioic Acid (C23:0) | 0.01 | 0.00 | 0.45 | 0.07 |
| 26. 1,24-Tetracosane Dioic Acid (C24:0) | 0.05 | 0.07 | 1.54 | 0.06 |
| <u>Hydroxylated 1,ω-Dicarboxylic Acids</u> | <u>0.18</u> | <u>0.02</u> | <u>1.19</u> | <u>0.12</u> |
| 27. (9)10,18-Dihydroxyoctadecenoic Acid<sup>1</sup> (C18:1) | 0.01 | 0.00 | 0.98 | 0.11 |
| 28. 9-Hydroxy-1,18-Octadecene Dioic Acid<sup>1</sup> (C18:1) | 0.17 | 0.02 | 0.21 | 0.02 |
| <u>1,ω-Diols</u> | <u>0.12</u> | <u>0.07</u> | <u>4.27</u> | <u>0.66</u> |
| 29. Hexadecane-1,16-diol (C16:0) |  |  | 0.27 | 0.03 |
| 30. Heptadecane-1,17-diol (C17:0) | 0.07 | 0.05 | 0.32 | 0.22 |
| 31. Octadecane-1,18-diol (C18:0) |  |  | 0.63 | 0.12 |
| 32. Eicosane-1,20-diol (C20:0) | 0.02 | 0.01 | 2.11 | 0.22 |
| 33. Docosane-1,22-diol (C22:0) | 0.03 | 0.01 | 0.94 | 0.07 |
| <u>Fatty Alcohols</u> | <u>10.59</u> | <u>0.89</u> | <u>18.33</u> | <u>2.04</u> |
| 34. Hexadecan-1-ol (C16:0) |  |  | 4.50 | 0.10 |
| 35. Heptadecan-1-ol (C17:0) |  |  | 2.35 | 0.11 |
| 36. Heptadecan-1-ol (C17:0)br |  |  | 0.53 | 0.06 |
| 37. Octadecen-1-ol (C18:1) |  |  | 3.13 | 0.15 |
| 38. Octadecan-1-ol (C18:0) | 2.95 | 0.25 | 4.30 | 0.19 |
| 39. Nonadecan-1-ol (C19:0) | 0.04 | 0.01 | 2.19 | 1.33 |
| 40. Eicosan-1-ol (C20:0) | 5.58 | 0.49 | 1.00 | 0.06 |
| 41. Docosan-1-ol (C22:0) | 2.02 | 0.14 | 0.33 | 0.04 |
| <u>Aromatics</u> | <u>3.30</u> | <u>0.32</u> | <u>4.22</u> | <u>1.01</u> |
| 42. Ferulic Acid | 1.55 | 0.14 | 3.01 | 0.38 |
| 43. Coumaric Acid | 1.75 | 0.17 | 1.21 | 0.64 |
|  | 100.00 | 8.72 | 100.00 | 9.01 |

Data are mean with SD ( $n = 4$ ). Only identified depolymerization products are reported in the table. <sup>1</sup>Photo-oxidation and auto-oxidation products.

In seeds and roots, the suberin monomer profiles were similar. A representative root suberin chromatogram is shown in **Figure 3B**. Small differences between these organs can mainly be attributed to the fact that seeds had a mixture of cutin and suberin monomers (Molina et al., 2008). To evaluate the contribution of embryo cutin to the total seed polyester composition, embryo and seed coat tissues were separated using a density gradient centrifugation method established for Arabidopsis (Perry and Wang, 2003). After transmethylation and GC-MS analysis of both fractions, it was determined that 50% of the HFAs, in particular the trihydroxy fatty acids, 18-hydroxy-octadecenoate, and DHPA are largely contributed by the embryo-enriched fraction (**Figure 4 A-B**) whereas 1,ω-diols, 1-alkanols (with exception of eicosan-1-ol), and dicarboxylic acids (DCAs; except for dimethyl octadecene-1,18-dioate) were mainly found in the seed coat-enriched samples (**Figure 4 A,C,D,E**). Similarly, polyester analyses of *B. napus* seed tissues showed that the trihydroxy 18:1 fatty acid fraction was largely found in embryo cutin, which also contained 18:1 and 18:2 DCAs, the major cutin monomers in this species (Molina et al., 2006). Although coumarate was exclusively found in the seed coat fraction, ferulate was present in both fractions, indicating that it is a common component of embryo in and seed coat lipid polyesters in *C. sativa* (**Figure 4 F)**. In spite of the high proportion of caffeate observed in leaf cutin, we did not detect this monomer in any of the seed fractions analyzed. Thus, in both *B. napus* and *C. sativa*, the embryo cutin composition seems to be different from that of leaf cutin with a predominance of 18:1 trihydroxy fatty acid. Conversely, most of the monomers that characterize suberin polyesters were located in the seed coat-enriched fraction.

**Figure 4:**
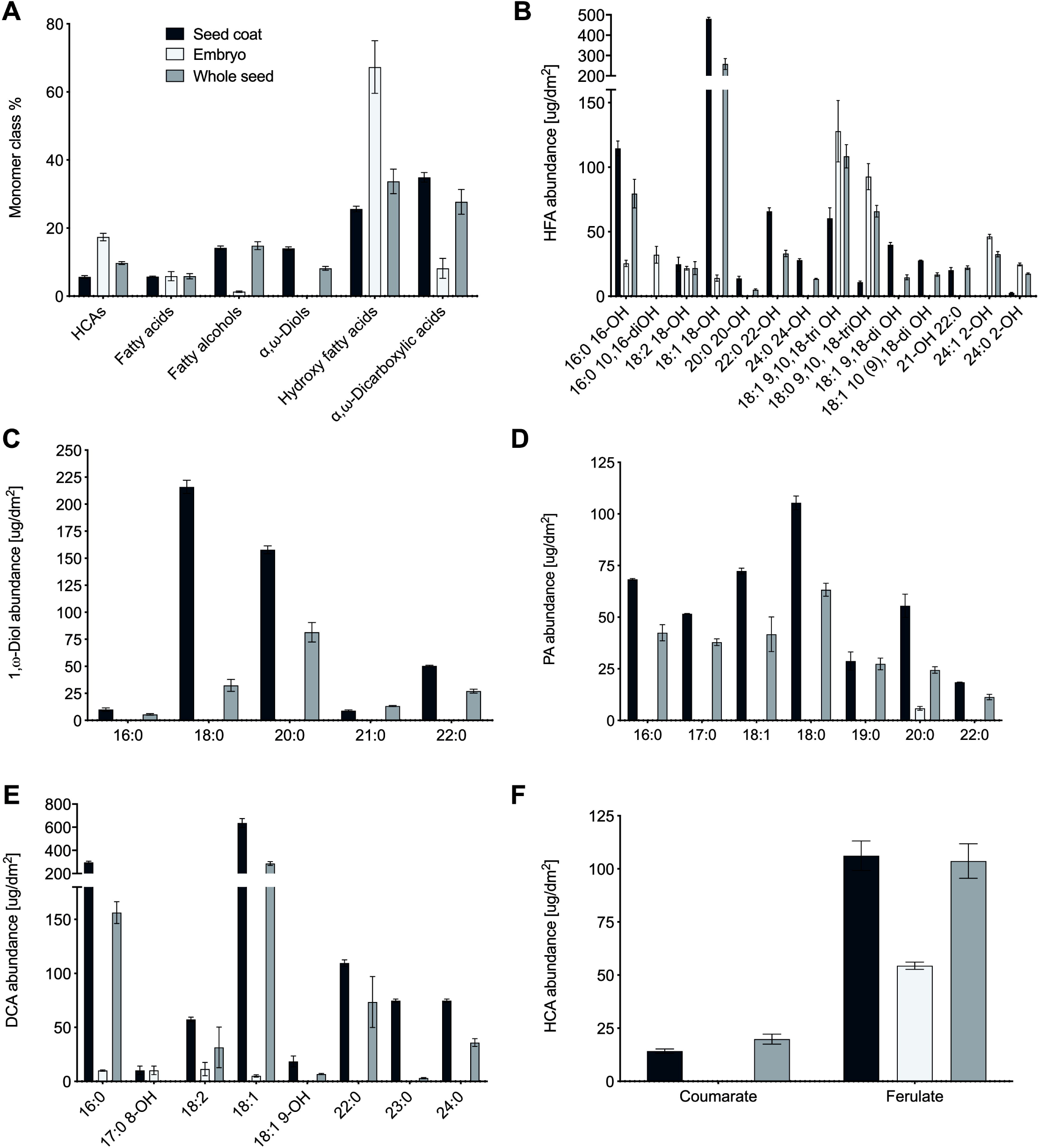
Lipid polyester monomer distribution in seed tissues. Comparison of transmethylation products from whole seeds, embryo-enriched and seed coat-enriched delipidated residues. (A) Relative content of cutin monomer classes. (B-G) Detailed seed coat, embryo and whole seed monomer composition in each component class, namely hydroxy fatty acids (HFA; B), 1,ω-Diols (C), primary alcohols (PA; D), dicarboxylic acids (DCA; E) and hydroxycinnamic acids (HCA; F). Error bars represent SE; *n*=3. Fatty acids did not present any particular distribution between seed tissues and are not included in this figure.

### 2.3. Lipid polyester monomer composition

#### Hydroxycinnamic acid derivatives

Although aromatics have been described as minor components of most cutins (Pollard et al., 2008), these monomers were unusually high in *C. sativa* cutin, constituting 28% and 23% of leaf and stem cuticular polyesters, respectively (**Table 2**). Notably, the most prominent hydroxycinnamic acid derivative in leaf cutin was caffeic acid (ca. 26% of all monomers), whereas coumaric and ferulic acids together accounted for 2% of the total monomers. Stem cutin, however, had similar proportions of coumarate, ferulate and caffeate. Transmethylation of flower residues yielded only aliphatic derivatives.

#### Dicarboxylic acids

*C. sativa* cutin contained saturated, unsaturated and mid-chain hydroxylated DCAs. Even-carbon number DCAs included 1,16-hexadecanedioate, 1,18-octadecanedioate, octadecene-1,18-dioate and octadecadiene-1,18-dioate, which were identified according to their retention times and by comparison to published mass spectra (Bonaventure et al., 2004; Holloway, 1982). In addition, a monomer with MS consistent with 18:3 DCA was observed in all three organs studied (**Supplemental Figure S1**). This may not necessarily be a α-linolenic acid derivative, given its late retention time relative to C18:1 and C18:2 DCAs (**Figure 2A**). Two saturated odd-chain DCA dimethyl esters (15:0 and 17:0) were identified by comparison to the MS of 15:0 DCA (Douglas et al., 1969), 16:0 and 18:0 DCA dimethyl esters (Eglinton and Hunneman, 1968; Holloway, 1982), and by their retention times. Small amounts of saturated even- and odd-chain DCAs ranging from C14 to C19 have been found in the algae *Botryococcus* (Douglas et al., 1969), and have been also reported in *Pinus radiata* stem cuticle (Franich and Volkman, 1982). Odd-chain aliphatics have also been described in cutin of other species. For example, apple fruit cuticles contain trace amounts of substituted, unsaturated C17 diacids, namely, methyl heptadecadiene-1,17-dioate and methyl heptadec-9-ene-1,17-dioate (Eglinton and Hunneman, 1968).

#### In-chain hydroxylated dicarboxylic acids

In addition to previously characterized 8(9)-hydroxy-1,18-octadecane dicarboxylic acid (Holloway, 1982), leaf cutin samples included three compounds tentatively identified as monosubstituted dimethyl esters, namely (6)7-hydroxy 15:0 DCA, 7(8)-hydroxy 16:0 DCA and 8-hydroxy 17:0 DCA. Two stereoisomer peaks with MS consistent with 7-hydroxy-C16:0 DCA were found, and the earlier peak overlapped with 18-hydroxy-oleate. Each peak contained a smaller proportion of the 8-hydroxy positional isomer. Their structure was also consistent with the mass spectra of the acetylated derivatives and their change in retention time compared to the trimethylsilyl derivatives (**Supplemental Figure 2**). Both 7- and 8-hydroxy 1,16-dicaboxylic acids have been also found in cutins of other angiosperm leaves, including *Coffea arabica, Auena satiua, Z. mays, Malus pumila, Citrus aurantifolia, Betula pendula, Encephalartos altensteinii, Pinus sylvestris*, and *Araucaria imbricata* (Holloway et al., 1972; Holloway and Deas, 1973; Hunneman and Eglinton, 1972). The compound (Figure 3A, peak #30) eluting about 0.2 equivalent chain length (ECL) after the *bis*-TMSi-dihydroxypalmitate peak (Figure 3A, peak #10), presented a dominant ion pair at 245/273 amu consistent with a mid-chain OTMSi-hydroxyl dicarboxylic acid dimethyl diester (8-HO-C17:0; Mr = 416), explaining the small m/z=401 peak (M-15). The mass spectrum of its acetylated derivative also supports this structure assignment (**Supplemental Figure 3**). The fact that a peak with MS consistent with 10,17-dihydroxy methylheptadecanoate (**Figure 3A**; peak #11) is found about 0.1 ECL after the putative 8-hydroxy dimethylheptadecandioate peak (**Figure 3A;** peak #30) makes this the most reasonable assumption. Similar retention behavior was observed between 16:0 7-hydroxy DCA and 16:0 10,16-dihydroxy FAME and between 18:0 9-hydroxy DCA and 18:0 10,18-dihydroxy FAMEs (**Figure 3A**). Minor amounts of substituted 17:0 DCAs have been also reported in depolymerized cutins of apple (Eglinton and Hunneman, 1968) and cranberry (Croteau and Fagerson, 1972).

Only flower cutin presented two stereoisomers tentatively identified 5,6-dihydroxy-15:0 DCA. Their mass spectra showed strong 273/203 ions, which could also correspond to a 5-OH C14:0 DCA (**Supplemental Figure S4**). Although retention times might vary with isomer position and are difficult to predict, as these compounds run very late, this peak more likely represents 5,6-dihydroxy-C15:0 DCA given the ion at m/z M-47. While this monomer has not been previously reported in cutins, methyl 8,9-dihydroxyheptadecane-1,17-dioate and methyl 9,10-dihydroxyoctadecane-1,18-dioate have been found in apple cuticles (Eglinton and Hunneman, 1968). We also detected trace amounts of the latter compound in leaf cutin transmethylation products. Furthermore, the monomer eluting before 5,6-dihydroxy-C15:0 DCA (peak #27, **Figure 3A**) seems to be structurally related, with a MS consistent with 6(7)-hydroxypentadecanedioate (Mr = 388), which in addition to the strong 273/217 and 259/231 pairs present peaks at m/z=373 (M-15), 257 (M-31) and 341 (M-47). (**Supplemental Figure S4**).

#### Substituted monocaboxylic acidsm

In addition to the highly represented (9)10,16-dihydroxy hexadecanotate monomers, small proportions of 10,17-dihydroxy heptadecanoate and (9)10(11),18-dihydroxy octadecanoate were identified in *C. sativa* leaf and stem cutin by comparison with the mass spectrum of (9)10(11),18-dihydroxyocatedecanoate (Holloway, 1982; **Supplemental Figure S5**). Additionally, these compounds had patterns of retention on a non-polar GC column consistent with these assigned identifications, with the mid-chain hydroxy dicarboxylic acid eluting immediately before the dihydroxy acid (**Figure 3A**). Two stereoisomers of 18:0 triol, characteristic of many cutins (Eglinton and Hunneman, 1968; García-Vallejo et al., 1997) also constituted minor components of the analyzed samples, along with 18:1 9,10-epoxy 18-hydroxy fatty acid methyl ester (identified by the characteristic methoxyhidrin MS; Holloway and Brown Deas, 1973).

Two-hydroxy fatty acids are often part of the depolymerization products from cutin- and suberin-containing tissues and were found in variable amounts in *C. sativa* leaf and stem cutin (7.8 and 11.8 Mol %, respectively, **Table 2**). However, their role as actual lipid polyester components is questionable; multiple lines of evidence implicate these as being derived from sphingolipids that are not fully removed by the delipidation process (Molina et al., 2006; Delude et al., 2016).

#### Photo-oxidation and auto-oxidation products

Another group of hydroxylated fatty acid derivatives was found in leaf cutin and suberin samples, and corresponded to monounsaturated dihydroxyoctadecanoic acids and monohydroxyoctadecanodioic acids. Leaf depolymerizates presented two monomers with m/z=285 as the base peak and had MS that agreed with described auto-oxidation products of the octadec-cis-9-ene-1,18-dioate (18:1 DCA) lipid polyester monomer (**Supplemental Figure S6A**). These corresponded to 8-hydroxy-cis/trans-9-ene products (m/z = 285), with only a small proportion of the 9-hydroxy-trans-10-ene (m/z = 271), and the *cis* (**Figure 3A**, peak # 33) eluting about 0.3 min ahead of the *trans* isomer (**Figure 3A**, peak # 34) in the hydrophobic column used in this work (Kosma et al., 2015). However, control assays with standards subjected to auto-oxidation conditions revealed that similar proportions of the products described above are synthesized (Kosma et al., 2015). Therefore, it is not entirely clear if the almost exclusive formation of 8-hydroxy-cis/trans-9-ene-1,18-dioate can be caused by non-enzymatic auto-oxidation or is a result of a novel enzymatic activity. On the other hand, root and seed suberin had predominantly the photo-oxidation product of 18:1 DCA, namely 9-hydroxy-trans-10-ene (m/z = 271), with a small proportion of the 8-hydroxy-cis/trans-9-ene products (m/z = 285), these last products being exclusively formed by auto-oxidation (**Supplemental Figure S6B**). Similarly, rutabaga periderm presented photo-oxidation derivatives of 18:1 DCA (Kosma et al., 2015).

*C. sativa* lipid polyesters also contained possible photo-oxidation and auto-oxidation products derived from 18-hydroxy-octadec-cis-9-eneoate (ω-OH-oleate). The major photo-oxidation products of this monomer are 10,18-dihydroxyoctadec-trans-8-enoic acid and 9,18-dihydroxyoctadec-trans-10-enoic acid, which yield m/z=271 and m/z= 315 diagnostic ions, respectively. Autoxidation gives similar proportions of 9,18-dihydroxy-trans-10-ene, 10,18-dihydroxy-trans-8-ene, 8,18-dihydroxy-cis/trans-9-ene and 11,18-dihydroxy-cis/trans-9-ene; the latter two isomers show m/z = 329 and 285 base peak ions and are not formed by photo-oxidation (Kosma et al., 2015). Based on the MS presented in **Supplemental Figure S6C**, *C. sativa* leaf tissues showed a mixture of isomers deriveded from both auto-oxidation and photo-oxidation of 18-hydroxyoleic acid. By comparison, a study performed in *Petroselium sativum* cutin demonstrates that photo-oxidation of unsaturated cutin components is the primary process occurring in this species (Rontani et al., 2005). *C. sativa* root and seed samples had predominance of the photo-oxidation derivatives of 18-hydroxyoleic acid (**Supplemental Figure S6D**), as reported for rutabaga periderm (Kosma et al., 2015).

## 3. Conclusions

Herein a comprehensive, qualitative and quantitative characterization of *C. sativa* apoplastic lipid polyesters has been provided, including those of leaf, stem, flower, seed coat and root. Commonly reported cutin and suberin monomers were detected. However, a number of novel cutin monomers were tentatively identified, which add to the complexity of polyester chemical profiles and the putative molecular conformations that these cutins can assume. This study further revealed striking differences between the lipid polyester monomer compositions of the aerial tissues of *C. sativa* and that of its relatives *Arabidopsis thaliana* and *Brassica napus* bringing to light questions on the validity of using cutin and suberin compositions for chemotaxonomic comparisons as well as evolutionary questions on the lineage specificity of lipid-based polyesters. Furthermore, it raises questions on yet to be identified biosynthetic enzymes and whether or not these enzymes are truly novel or are the result of neofunctionalizations of known enzymes that occur throughout the course of evolution and selective breeding. This may be particularly relevant given the hexaploid *C. sativa* genome.

Given the important functions of apoplastic polyesters in plant abiotic and biotic stress tolerance, these discoveries, in a species known for its inherent tolerance to abiotic stress, will help us to further comprehend the roles of different types of suberin and cutin polyesters in ameliorating the stressful conditions encountered in the field for yield maintenance.

## 4. Experimental

### 4.1 Plant material and growth conditions

Wild-type *Camelina sativa* (L.) Crantz cv. Celine seeds were obtained from Dr. Chaofu Lu (Montana State University, United States) and were surface sterilized in 70 % ethanol for 1 minute, in 50 % bleach for 5 minutes and washed 3 times with distilled water. The seeds were planted in PRO-MIX MPV potting mixture (Premier Tech Horticulture) and grown in the environmental chamber at 20-22 °C in an 18-h-light/6-h-dark photoperiod with 60–70 *µ*E m^-2^ s^-1^ light intensity and 65-75 % relative humidity.

### 4.2 Isolation of seed coat- and embryo-enriched fractions

Mature seeds (2.05g) were soaked in water for 24 h at 4 °C in darkness and further separated into embryos and seed coats using the method described in Perry and Wang (2003). Soaked seeds were gently crushed between two glass plates and placed into a Falcon tube (50 mL) with 5 mL of buffer MC (pH 7, 10 mM potassium phosphate, 0.5 mM sodium chlorate, and 0.1 M Sucrose). After centrifuging at 2000 rpm for 10 min, the supernatant was removed and the pellet resuspended in 25 % Percoll ^®^ (Sigma-Aldrich Canada, Oakville, Ontario) in buffer MC (12.5 mL Percoll^®^ and 37.5 mL buffer MC). The suspension was vortexed several times to release the remaining embryos and centrifuged at 1000 rpm at 10 °C to separate seed coats (supernatant) from embryos (pellet). The upper phase containing the seed coats and some embryos was removed and transferred to a new Falcon with 25 % Percoll^®^, vortexed and centrifuged; this procedure was repeated several times until seed coats and embryos were completely separated into two fractions. The isolated fractions were then washed with 5 volumes of 1 % (w/v) NaCl and stored at −80 °C.

### 4.3 Lipid polyester monomer preparation

For each replicate, seed coat fractions (isolated as detailed above), 100 mg of whole mature seeds, 20 partially or fully opened flowers, leaves # 14 to 17 (4 leaves in total), 4-week-old roots from plants grown on solid Murashige and Skoog Basal Medium, and 17 cm of the stem from the middle part of the plant (from the same stem region corresponding to that leaves # 14 to 17) were ground in liquid nitrogen using mortar and pestle. Following the protocol described in Jenkin et al. (2015), the samples were incubated in hot isopropanol at 85°C for 15 minutes, allowed to cool down, then further ground with a Polytron, and incubated for 24 hours at room temperature with a gentle shaking on a nutator, changing the solution to fresh isopropanol in between by spinning the tubes in the centrifuge at 1500 rpm for 1 minute, removing the liquid, and adding a fresh solution. The samples were sequentially extracted in each of the following solutions for 24 hours, changing the solution once to a fresh one in between: CHCl_3_:MeOH (2:1 v/v), CHCl_3_:MeOH (1:1 v/v), CHCl_3_:MeOH (1:2 v/v), and 100 % MeOH. All of the solution was removed, and the tubes were left open in the fume hood to dry for 2-4 days and then put in a desiccator containing anhydrous calcium sulfate to dry completely for approximately one week. Samples were subsequently depolymerized by NaOMe-catalyzed transmethylation, adding 1 mg g^-1^ DW each of methyl heptadecanoate (17:0 ME) and pentadecalactone as internal standards. The mixture was heated at 60°C for 2 hours with occasional vortexing. After cooling, methylene dichloride was added and the mixture was acidified with glacial acetic acid to obtain pH 4-5. The organic phase was washed twice with a dilute saline solution (0.5 M NaCl) and transferred to a clean tube. Anhydrous sodium sulfate was added to remove any remaining water, the tubes were centrifuged and the organic phase was transferred to a clean tube. After evaporating the solvent under a stream of nitrogen gas, the samples were derivatized by incubating in a mixture of 100 *µ*L of *N,O*-*bis*-(trimethylsilyl)-trifluoroacetamide (BSTFA) and 100 *µ*L of pyridine for 10 min at 110 °C. The derivatized samples were evaporated under a gentle stream of nitrogen gas, and re-suspended in heptane:toluene (1:1 v/v) for GC analysis.

### 4.4 GC-FID and GC-MS analysis

For monomer identification, representative samples from each tissue were analyzed on an Agilent 6850 gas chromatograph equipped with an Agilent 5975 mass spectrometer. Splitless injection was used with a DB5-MS column (30 m length, 0.25 mm inner diameter, 0.25 *µ*m film thickness). Temperature settings were as follows: inlet 350°C, detector 320°C, oven temperature program was set to 140°C for 3 min and increased to 310°C at a rate of 5°C per minute, oven temperature was then held at 310°C for 10 min. The helium flow rate was set at 1.5 mL per minute. The mass spectrometer was run in scan mode over 40-600 amu (electron impact ionization). For quantification, all replicates samples were analyzed on an Agilent 6890 gas chromatograph equipped with a flame ionization detector (FID). Split injection (20:1) was used with an HP5 column (30 m length, 0.25 mm inner diameter, 0.25 *µ*m film thickness). Temperature settings and helium gas flow rate were the same as those used for the GC-MS method. Peaks were quantified on the basis of their FID ion current. Peak areas (pA•sec) were converted to relative weights by applying FID theoretical correction factors, which assume that the FID response is proportional to carbon mass for all carbons bonded to at least one H-atom (Christie, 1991).

### 4.5 Light and fluorescence microscopy

Roots from 8-week-old plants were carefully brushed clean and thin sections were made by hand using a sharp double-edged razor blade. The sections were mounted in 50 % glycerol and observed under UV light using a using an Axio Imager M2 compound fluorescent microscope (Zeiss). Additional root sections were also stained with Sudan 7B solution (0.02 g of Sudan 7B in 5 ml of 95 % ethanol) and incubated at 70°C for 30 seconds and washed several times with distilled water. The stained sections were mounted in 50% (v/v) glycerol and observed under brightfield illumination.

### 4.6 Transmission electron microscopy

Root, leaf and stem samples (1 mm^2^ size), and whole seeds harvested at maturity (but that were not fully dry yet) were fixed with a mixture of 2.5% glutaraldehyde / 2% paraformaldehyde in 0.1 M cacodylate buffer and processed following the protocol detailed in Molina et al. (2009). Before the staining step, samples on grids were treated with 10% hydrogen peroxide for 10 min to enhance the contrast of both cutin and suberin (Heumann, 1990). Specimens were examined with a JEOL 100CX transmission electron microscope, and images processed with Adobe Photoshop CS2.

### 4.7 Scanning electron microscopy

Flower petals from 7-8 weeks old *C. sativa* plants were air dried on stubs and coated with gold-palladium particles using an Anatech Ltd Hummer VII sputter coater (Alexandria, Va). The samples were examined in a Vega\\XMU variable pressure scanning electron microscope (Tescan, Czech Republic) at an accelerating voltage of 15 kV.

## Acknowledgments

We thank Micaëla Chacón and Tanya Hiebert (Dept. of Biology, Carleton University) for assistance with the lipid extractions and Dr. Jianqun Wang (Carleton University Scanning Electron Facility) for assistance with the SEM imaging. We thank Prof. John Ohlrogge (Michigan State University) for use of gas chromatographs and general support. D.F. was supported by an undergraduate scholarship from the Brazilian government’s Science Without Borders Program. This work was funded by grants from the Natural Sciences and Engineering Research Council of Canada (I.M. and O.R), the USA National Science Foundation, grant #1547713 (D.K.K). This research was undertaken, in part, thanks to funding from the Canada Research Chairs program to I.M.

## Supplemental Figure Legends

**Supplemental Figure 1:** Mass spectrum of a molecule putatively identified as 18:3 1,18-dioate, dimethyl diester. The structure of this monomer was inferred by comparison to the mass spectra of 18:1 and 18:2 dicarboxylic acid dimethyl esters (Bonaventure et al., 2004). Although these are not strong diagnostic ions, as observed for dimethyl 1,18-octadeca-6,9-dienedioate (Christie, 2018), a small molecular ion is present, as well as [M−32]^+^ (*m/z*=304), [M−64]^+^ (*m/z*=272), [M−74]^+^ (*m/z*=262), and [M−92]^+^ (*m/z*=244), ions. The positions of the double bonds cannot be inferred from this mass spectrum.

**Supplemental Figure 2:** Identification of 8-hydroxy-hexadecane-1,16-dioate, dimethyl diester. Upper panel: Silylated derivative; lower panel: acetylated derivative. Comparison of acetyl and TMSi derivatizations indicate a single free hydroxyl group. The carboxyl end is reminiscent of 8- and 9-OTMSi cleavage of a FAME (m/z = 245, 259). This would either suggest that there are two isomers or that the molecule is a dicarboxylic acid. Adding up the Mr of fragments suggest the latter (245 + 259 – 102 = 402). Interpretation of m/z = 387 as (M-15) for TMSi derivative and m/z = 329 as (M – 43) for acetyl derivative. Fragmentations α to the -CH(OTMSi)-group at m/z = 245 and 259 place the OTMSi group at the 8-position for the major isomer, but there is a substantial amount of the 7-isomer (m/z = 231, 273). Acetylated mid-chain fragmentation pattern confirms assignment as (7)8-OH C16 DCA dimethyl diester. Subsequent analysis of retention times is also consistent with a mid-chain secondary OH group added to a DCA.

**Supplemental Figure 3:** Identification of 8-hydroxy-heptadecane-1,17-dioate, dimethyl diester. Upper panel: Silylated derivative; lower panel: acetylated derivative. The putative structure of this monomer was inferred from the fragmentation patterns of the two derivatives and their retention times.

**Supplemental Figure 4:** Preliminary identification of silylated pentadecanoate derivatives. A) Mass spectrum of 5,6-Dihydroxy-pentadecane-1,15-dioate, dimethyl diester. B) Mass spectrum 6(7)-hydroxy-pentadecanedioate, dimethyl diester (Mr = 388), which in addition to the strong 273/217 and 259/231 pairs present peaks at m/z=373 (M-15), 257 (M-31) and 341 (M-47). C) Mass spectrum of 9,15-dihydroxy pentadecanoate methyl ester.

**Supplemental Figure 5:** Mass spectra of 9,17-dihydroxy heptadecanoate (A) and (9)10(11),18-dihydroxy octadecanoate (B) were identified in *C. sativa* leaf and stem cutin by comparison to the mass spectrum of (9)10(11),18-dihydroxy octadecanoate (Holloway, 1982).

**Supplemental Figure 6:** Mass spectra of monomers potentially produced by photo-oxidation and auto-oxidation of fatty acids identified in leaf cutin (A, C) and suberin (B, D).

## Notes

### Competing Interest Statement

The authors have declared no competing interest.

