## Supplemental figures for "Extracellular lipids of *Camelina sativa*: Characterization of cutin and suberin reveals typical polyester monomers and novel functionalized dicarboxylic fatty acids"

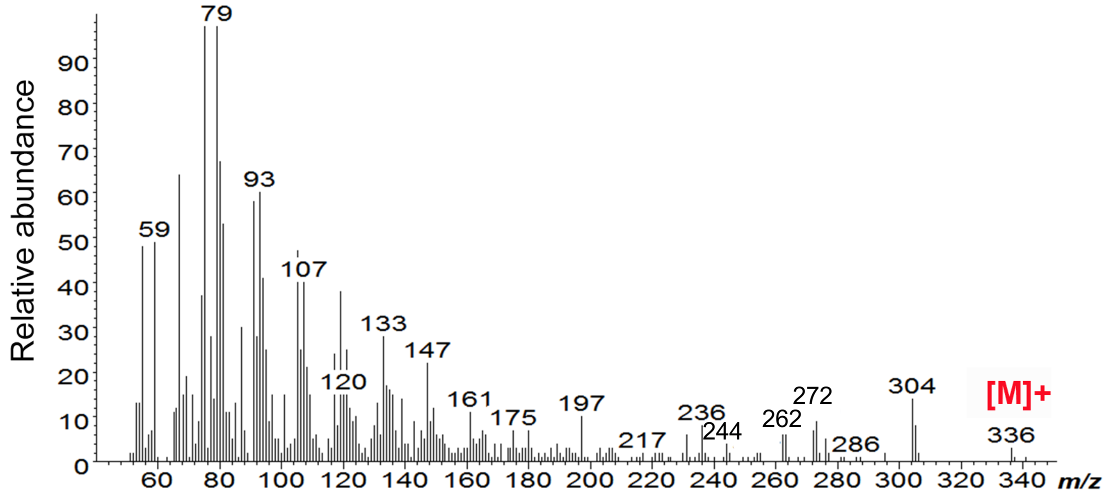


**Supplemental Figure 1.** Mass spectrum of a molecule putatively identified as 18:3 1,18-dioate, dimethyl diester. The structure of this monomer was inferred by comparison to the mass spectra of 18:1 and 18:2 dicaboxylic acid dimethyl esters (Bonaventure et al., 2004). Although there are not strong diagnostic ions, as observed for dimethyl 1,18-octadeca-6,9-dienedioate, a small molecular ion is present, as well as $[M-32]^+$ (*m/z*=304), $[M-64]^+$ (*m/z*=272), $[M-74]^+$ (*m/z*=262), and $[M-92]^+$ (*m/z*=244), ions. The positions of the double bonds cannot be inferred from this mass spectrum.

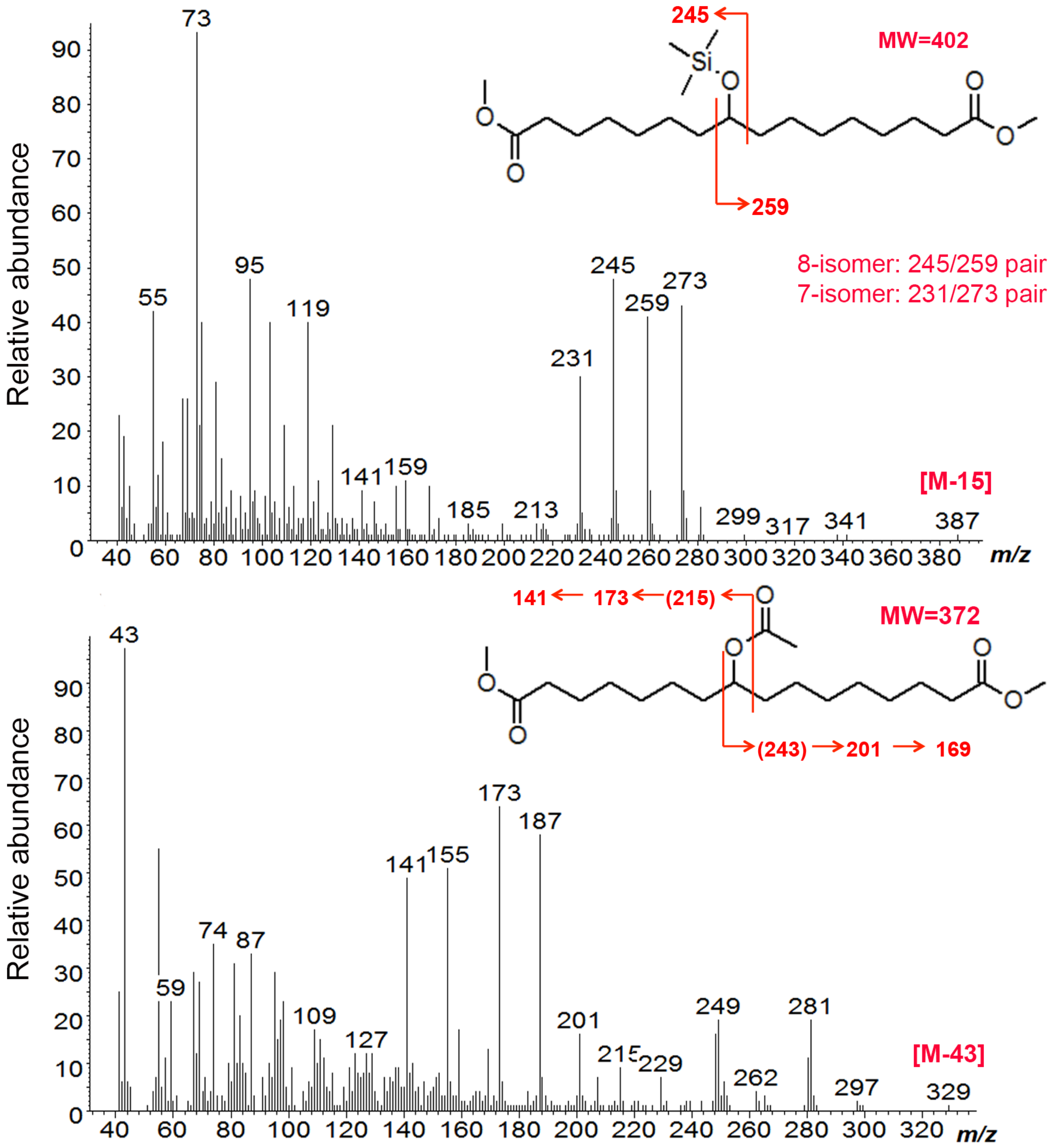


**Supplemental Figure 2.** Identification of 8-Hydroxy-hexadecane-1,16-dioate, dimethyl diester. Upper panel: Sylilated derivative; lower panel: acetylated derivative. Comparison of acetyl and TMSi derivatizations indicate a single free hydroxyl group. The carboxyl end is reminiscent of a 8- and 9-OTMSi cleavage of a FAME (m/z = 245, 259). This would either suggest that there are indeed two isomers or that the molecule can be a dicarboxylic acid. Adding up the Mr of fragments suggest the latter (245 + 259 – 102 = 402). Interpretation of m/z = 387 as (M-15) for TMSi derivative and m/z = 329 as (M – 43) for acetyl derivative fix the Mr of molecule. Fragmentations α to the -CH(OTMSi)- group at m/z = 245 and 259 place the OTMSi group at the 8-position for the major isomer, but there is a substantial amount of the 7-isomer (m/z = 231, 273). Acetylation mid-chain fragmentation pattern confirms assignment as (7)8-OH C16 DCA dimethyl diester. Subsequent analysis of retention times is also consistent with a mid-chain secondary OH group added to a DCA.

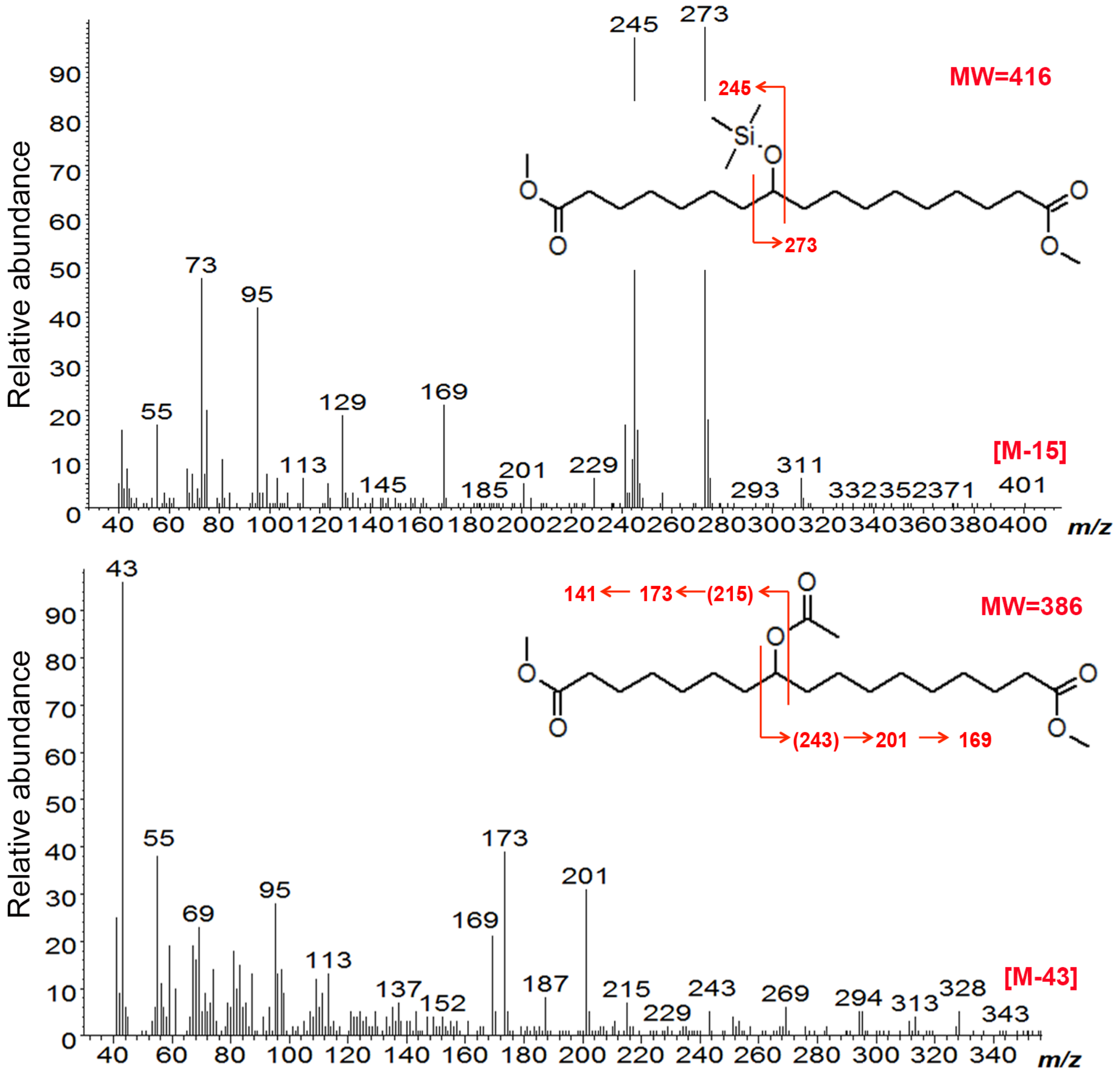


**Supplemental Figure 3.** Identification of 8-Hydroxy-heptadecane-1,17-dioate, dimethyl diester. Upper panel: Sylilated derivative; lower panel: acetylated derivative. The putative structure of this monomer was inferred from the fragmentation patterns of the two derivatives and their retention times.

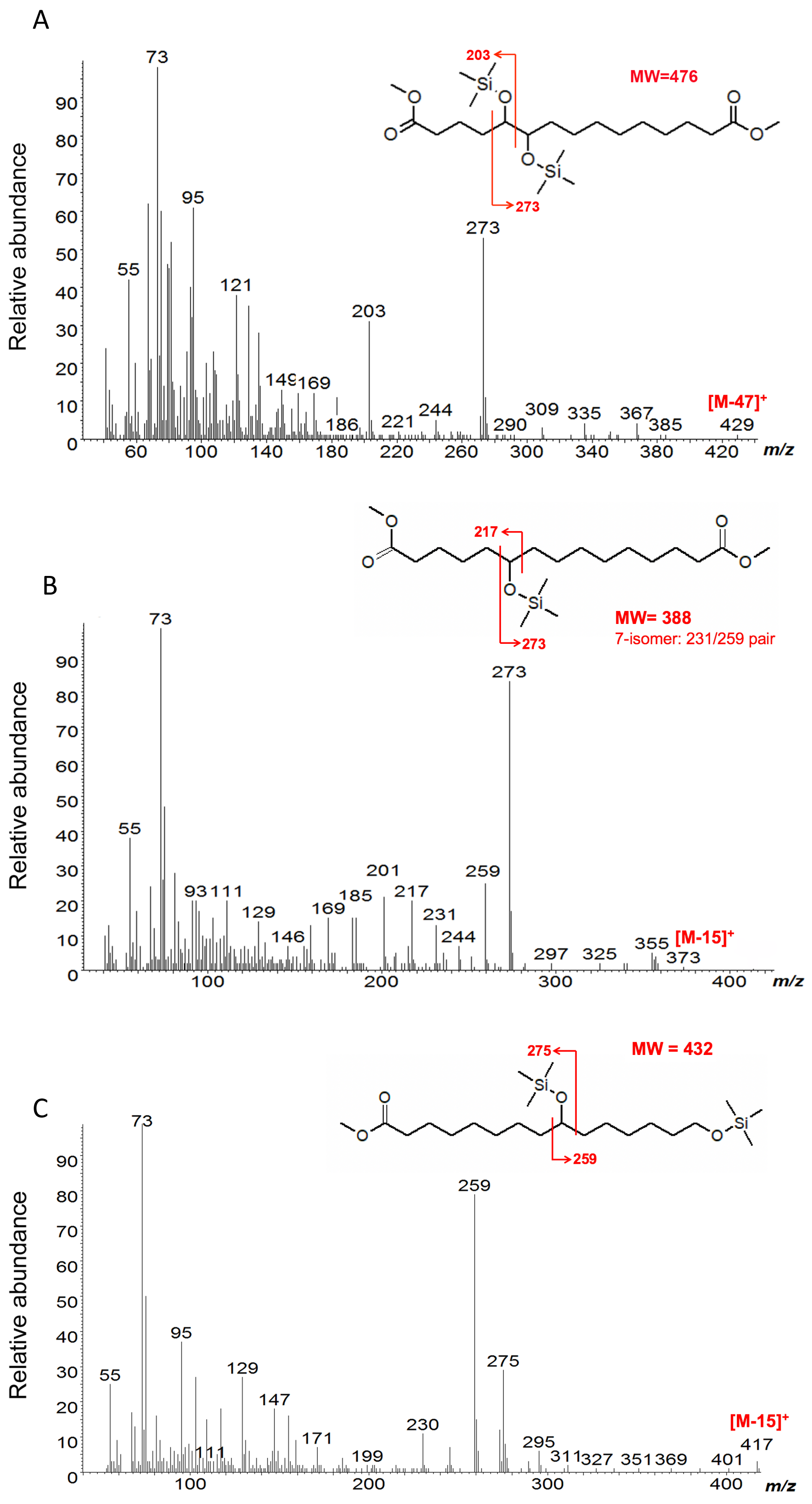


**Supplemental Figure 4.** Preliminary identification of sylilated pentadecanoate derivatives. A) Mass spectrum of 5,6-Dihydroxy-pentadecane-1,15-dioate, dimethyl diester. B) Mass spectrum 6(7)-hydroxy-pentadecanedioate, dimethyl diester (Mr = 388), which in addition to the strong 273/217 and 259/231 pairs present peaks at m/z=373 (M-15), 257 (M-31) and 341 (M-47). C) Mass spectrum of 9,15-dihydroxy pentadecanoate methyl ester.

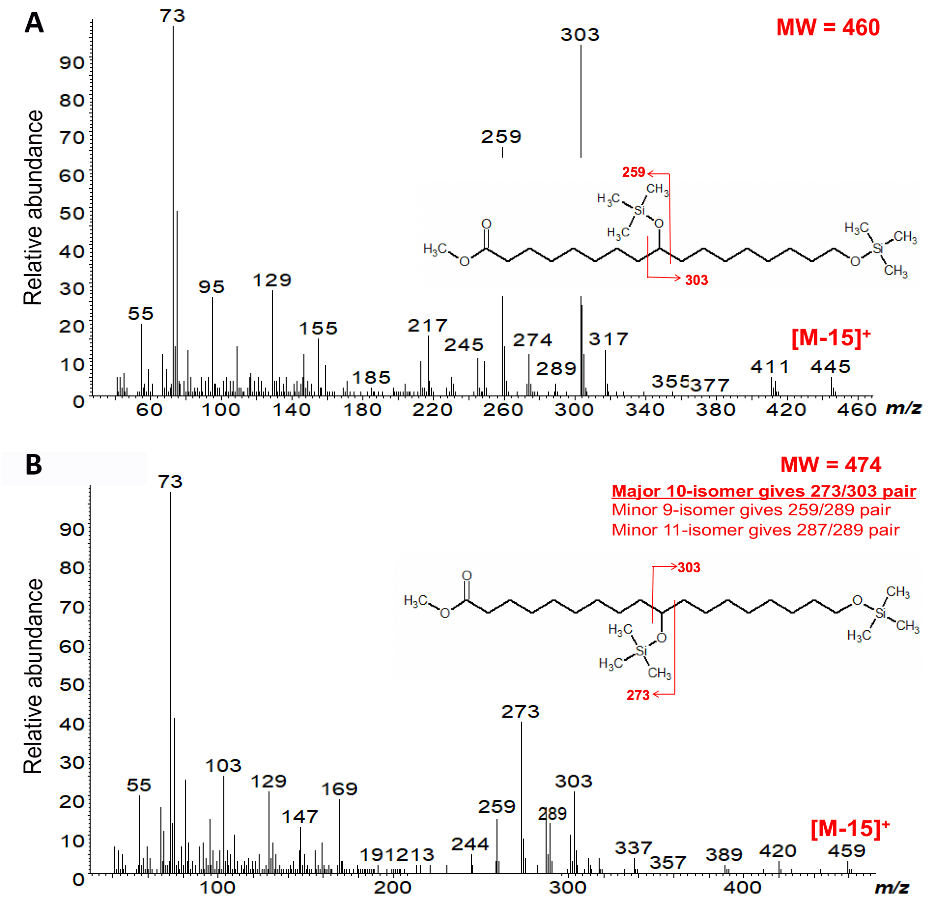


**Supplemental Figure 5.** Mass spectra of 9,17-dihydroxy heptadecanoate (A) and (9)10(11),18-dihydroxy ocatedecanoate (B) were identified in *C. sativa* leaf and stem cutin by comparison to the mass spectrum of (9)10(11),18-dihydroxy ocatedecanoate (Holloway, 1982).

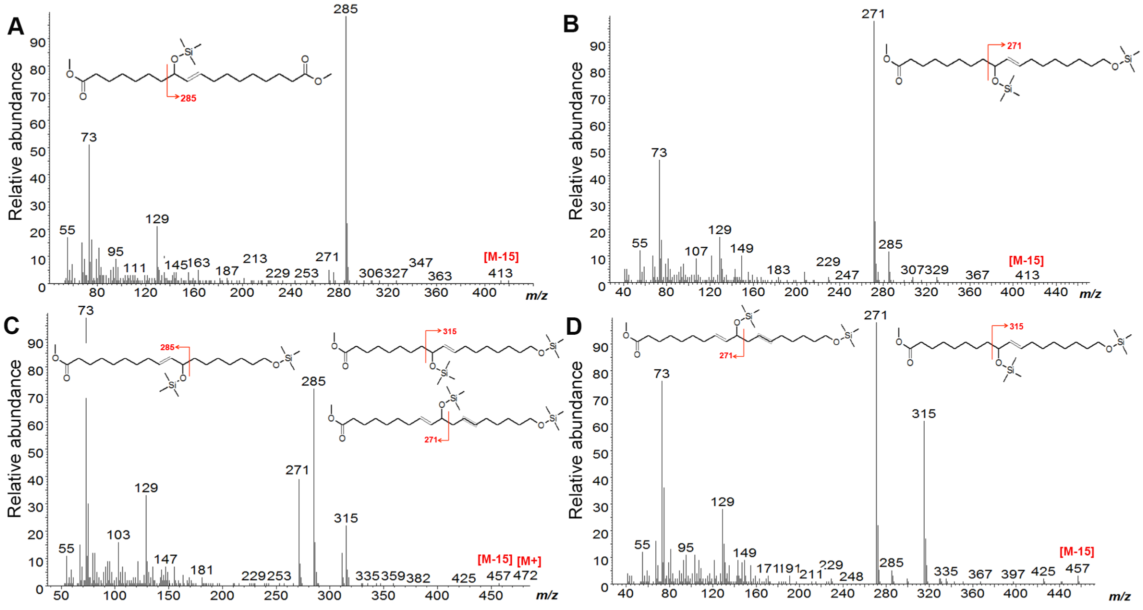


**Supplemental Figure 6.** GC-MS (EI) analysis of monomers potentially produced by photo-oxidation and auto-oxidation of fatty acids identified in leaf cutin (A, C) and suberin (B, D).
